# Active inference, stressors, and psychological trauma: A neuroethological model of (mal)adaptive explore-exploit dynamics in ecological context

**DOI:** 10.1101/695445

**Authors:** Adam Linson, Thomas Parr, Karl J. Friston

## Abstract

This paper offers a formal account of emotional inference and stress-related behaviour, using the notion of active inference. We formulate responses to stressful scenarios in terms of Bayesian belief-updating and subsequent policy selection; namely, planning as (active) inference. Using a minimal model of how creatures or subjects account for their sensations (and subsequent action), we deconstruct the sequences of belief updating and behaviour that underwrite stress-related responses – and simulate the aberrant responses of the sort seen in post-traumatic stress disorder (PTSD). Crucially, the model used for belief-updating generates predictions in multiple (exteroceptive, proprioceptive and interoceptive) modalities, to provide an integrated account of evidence accumulation and multimodal integration that has consequences for both motor and autonomic responses. The ensuing phenomenology speaks to many constructs in the ecological and clinical literature on stress, which we unpack with reference to simulated inference processes and accompanying neuronal responses. A key insight afforded by this formal approach rests on the trade-off between the epistemic affordance of certain cues (that resolve uncertainty about states of affairs in the environment) and the consequences of epistemic foraging (that may be in conflict with the instrumental or pragmatic value of ‘fleeing’ or ‘freezing’). Starting from first principles, we show how this trade-off is nuanced by prior (subpersonal) beliefs about the outcomes of behaviour – beliefs that, when held with unduly high precision, can lead to (Bayes optimal) responses that closely resemble PTSD.

## 1. Introduction

This paper presents software simulations that extend previous theoretical work on a computational psychiatry model of PTSD (Linson and Friston, in review). The simulations situate this work in relation to computational ethology (Anderson and Perona, 2014) and neuroeconomics (Watson and Platt, 2008). We use a Markov decision process model and active inference to present some scenarios – and sequential responses – that this minimal model can exhibit, on the view that, with the appropriate parameterisation, other natural behaviours can be emulated (Metz et al., 1983). With this approach, neuroethological and ecological constructs are shown to emerge from first principles (Daunizeau, 2018; Linson et al., 2018; Ramstead et al., 2018).

After demonstrating the basic architecture and ensuing active inference, we then consider the emergence of adaptive and maladaptive (pathological) responses to various cues in terms of constructs related to threat and fear. We then connect these constructs to the symptomology of PTSD, to provide a formal account of its pathophysiology. Specifically, by formulating a generative model under active inference, we can trace the implied message passing to its neurobiological substrate. Connecting the generative model to computational neuroanatomy ensures that one can predict empirical responses as measured using (e.g.) electrophysiological and neuroimaging data, for eventual translation into clinical and therapeutic applications (Parr and Friston, 2018).

This paper comprises two parts. In the first, we briefly overview active inference and the particular model used to deconstruct emotional inference in stressful situations. This model is presented in some detail, from first principles, using minimal but plausible assumptions about embodied inference. We will spend some time demonstrating the emergent phenomenology – both in terms of belief updating and the accompanying responses at both the neuronal and behavioural (motor and autonomic) levels. The second part of this paper then revisits these simulations in light of established constructs and phenomena in the literature on stress-related behaviour; with a special focus on pathology of the kind associated with post-traumatic stress disorder (PTSD).

## 2. Methods

We start by describing a generative model of active engagement with a simple world. Crucially, this model is specified in terms of biologically plausible contingencies and straightforward physical laws, without any reference to valenced states or emotional constructs. The agenda here is to demonstrate how emotional behaviour emerges from inference based upon sensory evidence that, in itself, has no particular valence or meaning. Another important aspect of this formulation of ‘emotional inference’ is its multimodal nature; namely, the perceptual synthesis or inference based on exteroceptive, proprioceptive and – importantly – interoceptive sensory cues. This means the model necessarily generates sensory outcomes in multiple domains, such that model inversion constitutes multisensory integration in the service of informing emotional behaviour. In active inference, policy selection and adaptive behaviour is treated as an inference problem, much in the spirit of planning as inference (Attias, 2003; Baker et al., 2009; Botvinick and Toussaint, 2012; Kaplan and Friston, 2018).

Formally, active inference under (partially observed) Markov decision process models requires a specification of hidden states and their sensory consequences (Friston et al., 2017a; Friston et al., 2017b). Once the hidden states and outcomes have been established, the parameters of the generative model determine the likelihood a particular outcome is generated by a combination of hidden states. This is usually encoded in an **A** matrix. The transitions among hidden states are parameterised in terms of **B** matrices (for each hidden state). Each hidden state is equipped with probability transition (**B**) matrices that are controlled by action, so that there is a repertoire of actions for each hidden state. Prior preferences – that determine the quality or expected free energy of allowable policies – are encoded by a **C** vector over each outcome modality. Finally, beliefs about initial states are specified with **D** in a matrix for each hidden state. Equipped with this model specification, one can then use standard variational procedures to simulate active inference in terms of belief updating and subsequent policy selection (i.e., behaviour); for details, see Friston et al. (2017b).

A key aspect of this belief updating is that agents entertain (posterior) beliefs about states of the world (including the body) *and their action upon the world*; namely, the policies that determine state transitions. Recognising or inferring states of the world proceeds using standard Bayesian observer assumptions. Technically, this involves minimising a *variational free energy* (upper) bound on surprise (a.k.a. self-information). This is mathematically equivalent to maximising a (lower) bound on model evidence (a k.a. self-evidencing) (Hohwy, 2016; Winn and Bishop, 2005). This optimisation can be formulated in a biologically plausible way by associating neuronal dynamics with a gradient flow on variational free energy (Friston et al., 2017a). The special aspect of active inference pertains to policy selection based upon inferences about which policy is being pursued (from which the next action is selected). Crucially, these inferences are based upon prior beliefs that policies minimise *expected free energy*. Minimising expected free energy can be conceptualised as selecting policies that minimise uncertainty, while leading to preferred (i.e., expected) outcomes. In other words, expected free energy combines epistemic and instrumental imperatives in a way that dissolves the exploration-exploitation dilemma (Friston et al., 2015b).

The particular generative model used in this paper – to demonstrate the emergence of healthy and pathological responses to safe and unsafe circumstances – aims to be as simple as possible, while being sufficiently comprehensive to explain the phenomena of interest; namely, (neuro)ethological constructs related to threat and fear, and the symptomology and pathophysiology of PTSD (Linson and Friston, in review). To (literally) cartoon the structure of this generative model, we will use a ‘Tom and Jerry’ analogy, bearing in mind that exactly the same probabilistic structure can be applied to any situation involving defensive responses to existential integrity; ranging from the pathology of predation through to psychosocial interactions.

The metaphor we have in mind considers a mouse (Jerry), who must decide how to behave to avoid a cat (Tom). To set up Jerry’s generative model of his environment, we start by outlining the sensory modalities that this model must account for. Jerry’s outcome modalities cover all sensory domains relevant for his inference about states of the world and policies (Figure 1). His exteroceptive modalities are *auditory* and *visual*. The auditory modality has three levels (*silence, soft sounds* and *loud sounds*), and the visual modality has five levels (an *empty horizon*, a *dot on a horizon*, a *cat shape*, a *dog shape*, and a *blur*). In addition, he has a *proprioceptive* modality with a sensation that signals (self) *movement* or *not*, and a *baroreceptive* modality that repo1ts bodily pulsations that are either *palpable* or *not* (con-esponding to high or moderate-to-low BPM). The latter two modalities are jointly specified in four combinations. The implicit hypotheses Jerry may appeal to (sub-personally) to explain these senso1y data are separated out into three sorts (factors) of hidden state. The first is the *other creatures* in his vicinity, which takes five possible values: the absence of other creatures, and either Tom or Spike are near or far away, respectively. The second is whether Jeny himself is *moving*, which he may or may not be, and the third is his own *heart rate*, which may or may not be accelerated.

**Figure 1.**
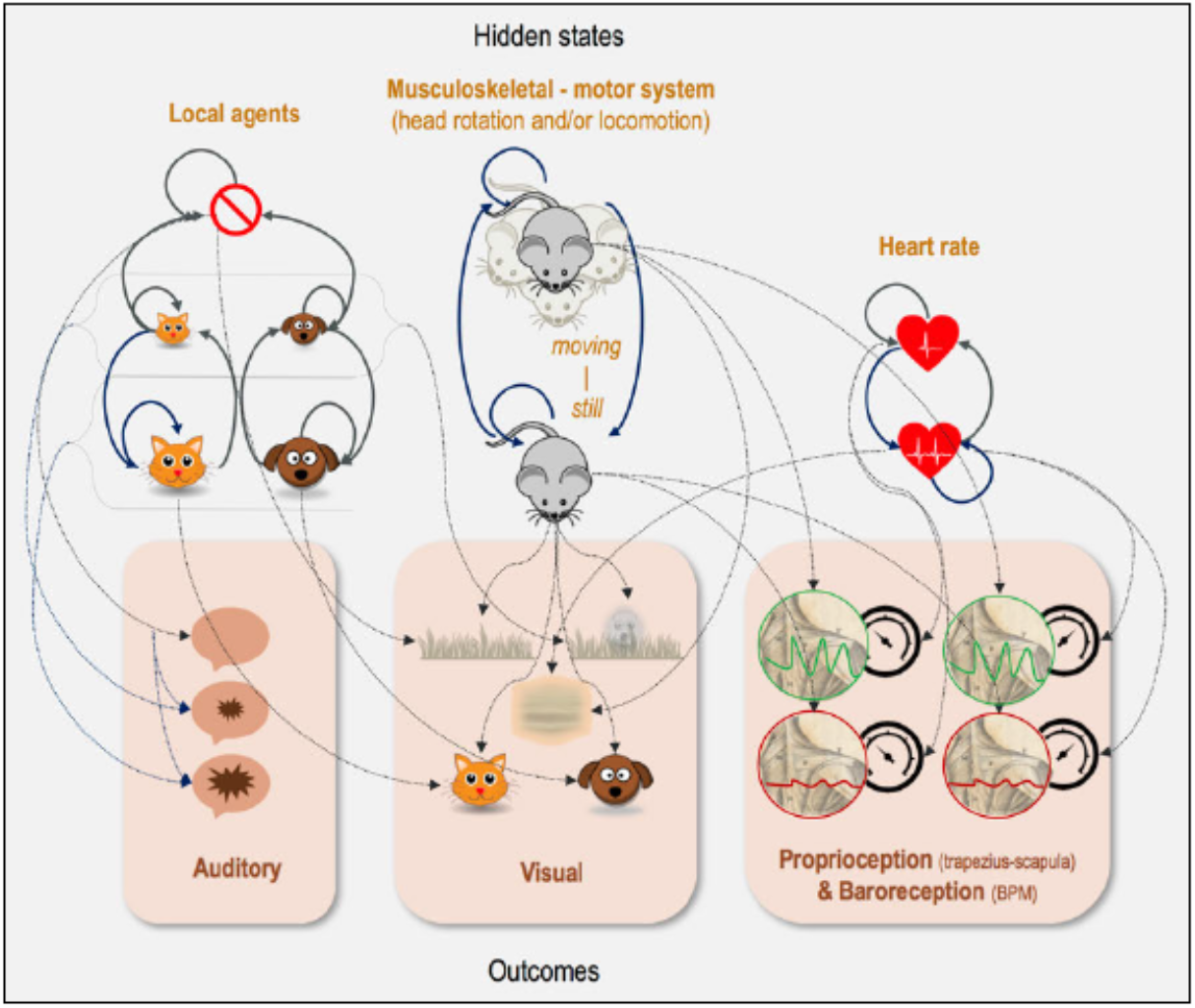
The generative model. This schematic shows the form of the generative model we assume Jerry uses to make sense of his environment. The hidden states represent the hypotheses he may appeal to in order to explain the sensory outcomes he experiences. The mrows indicate conditional probabilities, such that an arrow from one pictogram to another indicates the dependence of the second on the first. Please see the main text for details of this model. Brief summary – environmental states (’ local agents’ icons): no agents, distal cat, distal dog, proximal cat, proximal dog; musculoskeletal-motor system states (see icon labels): moving, still; cardiac states (top to bottom): moderate-to-low, high; auditory outcomes (top to bottom: silence, soft sounds, loud sounds; visual outcomes (see icons): empty horizon, dot on horizon, cat shape, dog shape, blur; proprioceptive outcomes (see icons): moving or not; baroreceptive outcomes (see icons): palpable (high BPM) or not (moderate-to-low BPM).

With these factors and modalities in place, we can now consider the likelihood mapping (A) that describes how Jeny (implicitly) believes these hidden state factors give rise to their associated outcomes, and the action-dependent transitions among hidden states that underwrite beliefs about self and environmental trajectories. The likelihood of auditory outcomes depends upon the *other creatures* in Jerry’s vicinity. The closer either Tom or Spike are, the louder the sounds Jerry can expect to hear. On rare occasions, a soft or loud sound may occur in the distance; even when no agents are present (e.g., a tree falling). For the visual outcome, proximal cats or dogs give rise to clear visual cat or dog forms, respectively. If they are further away, both give rise to an indistinct form on the horizon, and if they are absent, the horizon is empty. In addition, the visual outcome depends upon whether or not Jerry is moving and his heart rate. If he is moving with a high heart rate, all Jerry sees is a blur, regardless of who is in his vicinity. If he is moving without a high heart rate and there are distal creatures present, it is equiprobable that he will see a dot on the horizon or an empty horizon. Proprioceptive and baroreceptive data report, respectively, whether or not Jerry is moving, and whether or not his heart rate is rapid.

Next, we specify the dynamic structure or contingencies of the generative model: Jerry may engage in four different sorts of action, grouped into policies (sequences of actions), which have implications for all three hidden state factors. The actions that comprise these policies include: roaming/scanning; orienting; fight-or-flight; and freezing. There are seven policies (Figure 2), three of which involve fight-or-flight actions, three of which involve freeze actions, and one which involves neither fight-or-flight nor freeze. All policies begin with roaming/scanning followed by orienting. Policies 1 and 4 continue with an unrelenting fight-or-flight or freeze, respectively. For each of these, there is a corresponding policy that turns into orienting (2 and 5), and one that reverts to roaming/scanning (3 and 6). The final policy (7) corresponds to an alternating roaming/scanning and orienting policy (see Figure 2b).

**Figure 2.**
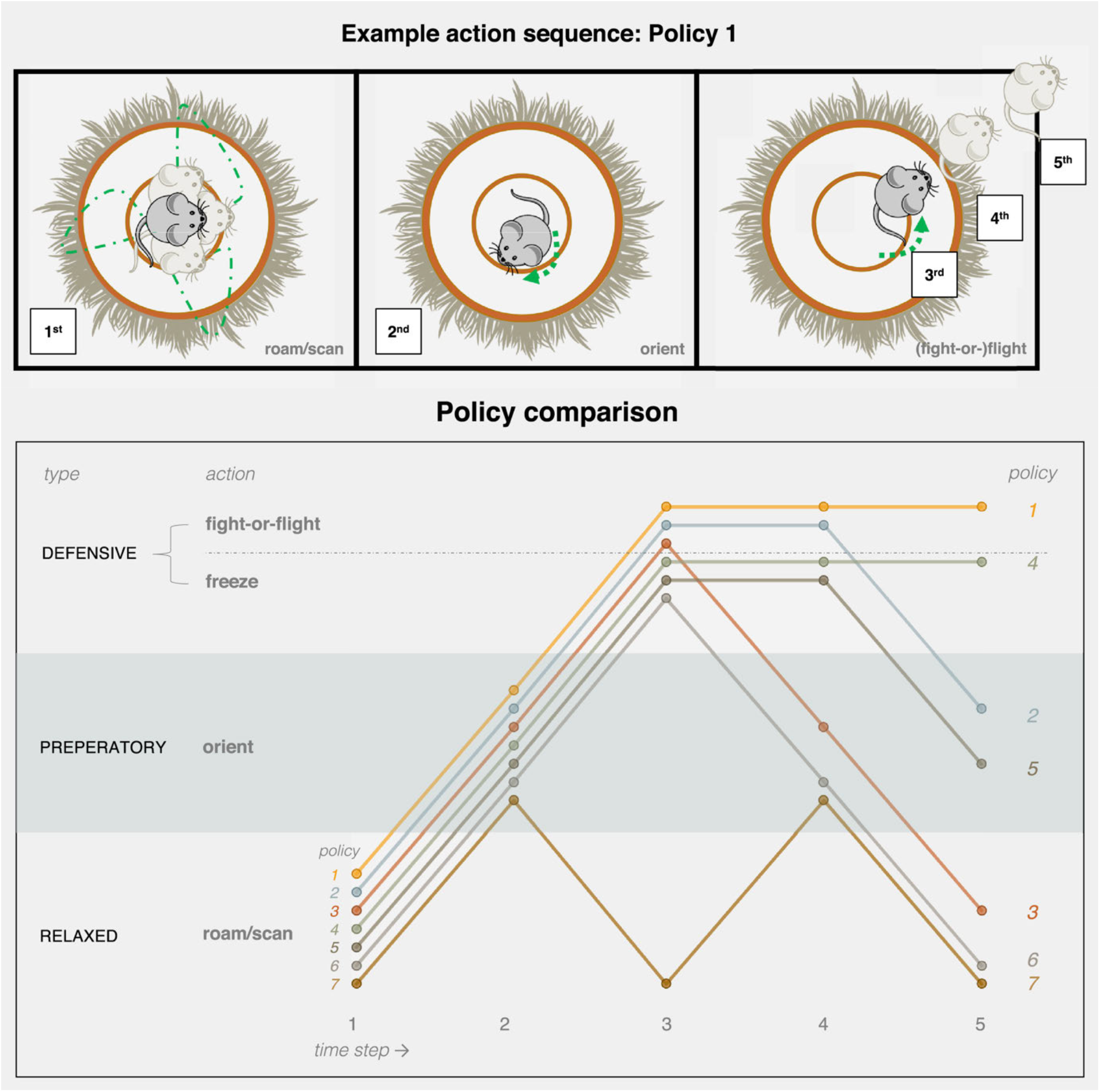
Policies. This schematic illustrates the alternative plans (or sequences of actions) that Jerry may choose to pursue. The five actions of Policy 1 are cartooned in the upper plot, which shows the following sequence: Jerry begins by (1) roaming/scanning his environment, continues on to (2) orienting, and finally, he engages in (3-5) a series of fight-or-flight actions. This sequence can be compared with the other six policies, indicated in the lower plot. To interpret this schematic, note the labels on the left that specify each action (which have been grouped into action types in the far left column). Each line is an alternative policy, with circles indicating the action at each time-step. The schematic acts as a key to interpret the labelling of the 7 alternative policies available to Jerry. This policy numbering will be adopted consistently in subsequent figures.

These actions contextualise the transition probabilities associated with each hidden state factor. The first transition (**B**) matrix allows the creatures in Jerry’s vicinity to change their location according to the arrows shown in Figure 1. For example, a nearby cat can transition probabilistically to a distant cat, but cannot change into a nearby dog. This transition depends on whether Jerry engages in his sympathetic *fight-or-flight* response, which places Jerry farther from Tom, or his parasympathetic *freeze* response, which places Tom farther from Jerry. These policy dependent contingencies manifest as Tom moving away. Jerry can similarly get away from Spike. However, when Spike wanders into the vicinity, Tom stays away. The other two policies lead to a greater probability that Tom approaches, attracted by Jerry’s movement or heart rate. With respect to Jerry’s intrinsic hidden states, when engaging either the *roaming/scanning* or *fight-or-flight* actions, he transitions into moving, and otherwise, if *orienting* or *freezing*, he transitions to still. When engaged in *fight-or-flight* or *orienting*, his heart rate becomes rapid, which is counteracted by the remaining two options.

Having specified the way in which data are generated from changing states, we can now specify Jerry’s prior preferences (**C**). He slightly disprefers seeing a blur, moderately disprefers seeing a dot on the horizon, and strongly disprefers seeing the outline of his feline predator (Tom). But, he is happy to see an empty horizon or the outline of a dog (Spike), which means Tom won’t be there. In the auditory domain, Jerry prefers not to hear any sound, slightly disprefers soft sounds, and strongly disprefers loud sounds. In the jointly specified proprioceptive and baroreceptive domains, he prefers to be relaxed and moving (the result of roaming/scanning), is neutral about being excited and still (due to orienting), but disprefers the remaining combinations (the result of fight-or-flight or freezing). Note in using loaded terms like prefer and disprefer, we simply mean that the generative model includes prior probabilities about the kinds of outcomes the agent expects to encounter *a priori*.

All that remains is to set up prior beliefs about the initial states (**D**), which provides two alternative simulation scenarios (see Results). In the first, Jerry assumes he is in a safe location (‘cat poor’), represented by flat priors. In the second, he is in an unsafe location (‘cat rich’), represented by a strong prior belief that Tom is approaching from a distance. We will see below the key differences in how these scenarios affect Jerry’s neurobiology and behaviour.

### 2.1 Biological grounding

Certain aspects of this generative model are motivated by known physiology. For example, policies (action sequences) are defined such that orienting always occurs before and after engaging in ‘defensive’ actions; namely, fight-or-flight and freezing. Orienting entails holding still, which increases visual acuity (e.g., following a head rotation to make a sound source visible). It also entails cardioacceleration (along with vasodilation in skeletal muscle) to increase motor readiness. Physiologically, this corresponds to noradrenergic arousal and vagal and/or sympathetic activation (Berridge, 2008; Dampney, 2019; Hilton, 1982), which has been found to be enhanced in PTSD (Ronzoni et al., 2016).

The implications of these contingencies can be summarised as follows. When Jerry is roaming/scanning, his proprio- and baroreceptive outcomes are as he prefers (i.e., expects *a priori*). And yet, this also means he is too relaxed to adequately respond to an event of interest. Moreover, it is possible Tom is on the horizon, in or out of focal range. Thus, he must intermittently orient to prepare for motor action and subsequent epistemic foraging (which may include turning toward a distal sound source, to bring it into his visual field). Similarly, if he is moving rapidly in fight-or-flight, a blurry outcome ensues, as a proxy for degraded visual acuity – a fundamental consequence of biophysics from insects to mammals (Land, 1999). Holding still is a countermeasure, so that Jerry can assess whether or not it is safe to lower his defences (i.e. reliably resampling the scene to accumulate evidence for his posterior beliefs).

The proprioceptive outcomes were selected as a reasonably economical way to infer self-movement pertaining to forelimb use (Roberts, 1974) and/or head rotation (DeToledo and David, 2001), active in roaming/scanning, and fight-or-flight, and inactive in the remaining two policies (as orienting is defined here by a contingent completed head turn, in contrast to stationary scanning). These outcomes originate from a particular head-trunk interface site found across species; homologous to the trapezius/scapula attachment in humans. This locus of proprioception is used here for neuroethological generalisation (see Discussion), given its extensive evolutionary history from 400 million year old fish up to present-day mammals (Matsuoka et al., 2005).

In short, this minimal generative model is equipped with everything we need to account for, in terms of adaptive behaviour that has consequences in the exteroceptive, proprioceptive, and interoceptive domains. In the following, we show how inversion of this model – using standard Bayesian message passing schemes – gives rise to sensible behaviours under the prior preferences and beliefs outlined above. We then repeat the simulations using aberrant priors to illustrate the emergence of maladaptive behaviour – in a way that reproduces the symptomology of PTSD, and in turn points to its possible pathophysiology (see Linson and Friston, in review). As noted above, once the generative model is specified, the requisite (planning as) active inference can be simulated using standard variational (marginal) message passing, whose neurobiological plausibility has been established to a certain degree in several settings: see (Friston et al., 2017a) for an introduction to the neuronal process theories assume in what follows.

## 3. Results

To build an intuition as to how active inference works under this sort of model, we start by describing some simple scenarios. While there are many scenarios and response sequences that this simple model can exhibit, we focus on two main narratives that best illustrate the relevant inference and behaviour. We first introduce the two scenarios and continue by describing four variants that characterise our analysis.

In Scenario 1 (Figure 3a), imagine Jerry is roaming/scanning, which he infers via proprioceptive movement cues and impalpable baroreceptive pulsation. For the initial exteroceptive outcomes, he sees an empty horizon, but hears a soft sound. Given his beliefs about what’s ‘out there’, he considers there to be a small chance that no agents are present, and a good chance that either Tom or Spike are approaching. The concurrent auditory and visual outcomes lead him to orient, allowing for some uncertainty resolution about the hidden state via the subsequently foraged visual outcome. If he then sees an empty horizon or close-up dog shape, he infers he can safely resume roaming/scanning. However, if he sees a close-up cat shape, it becomes immediately clear that a defensive policy is mandated.

**Figure 3.**
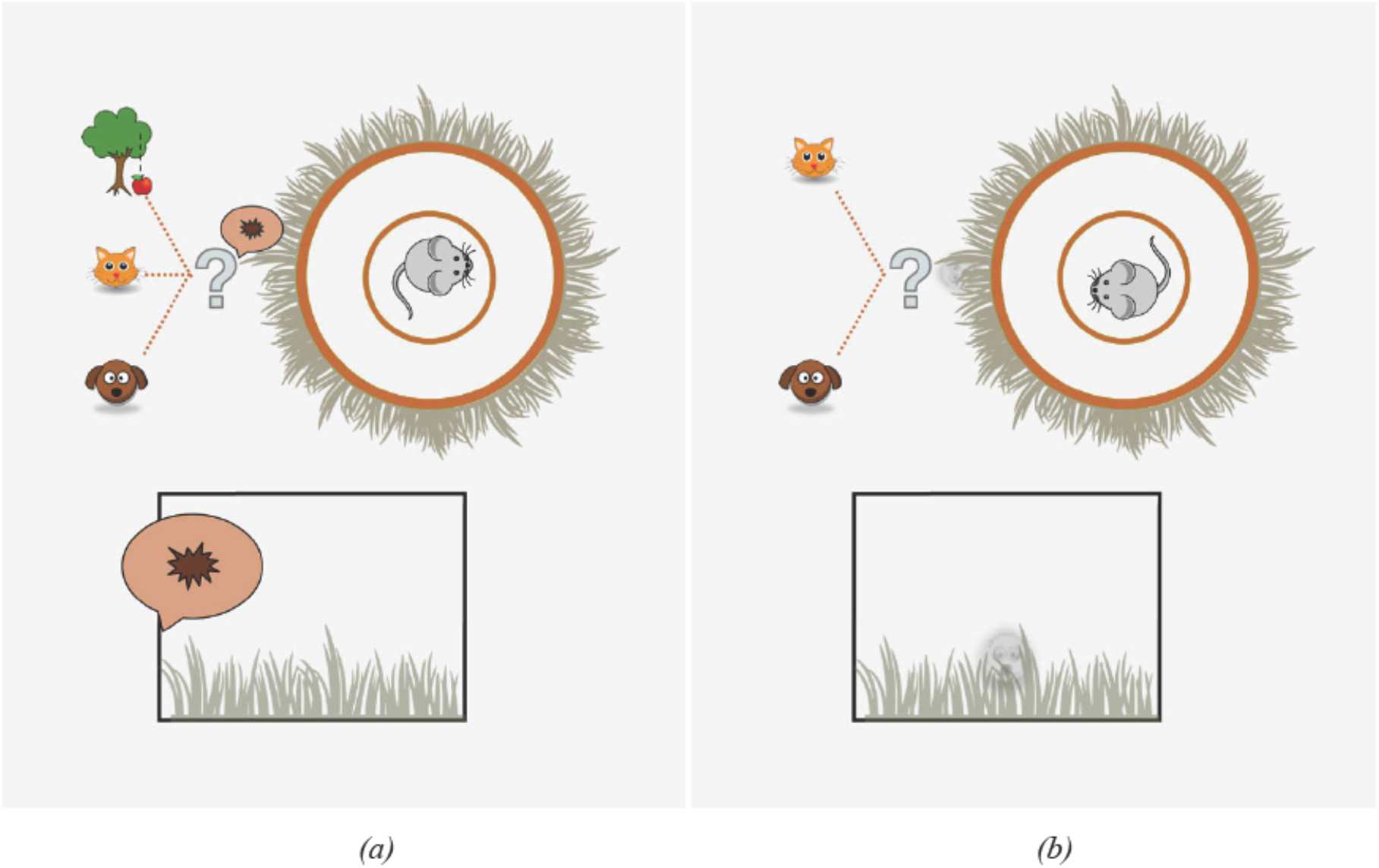
Scenario 1 (a), Scenario 2 (b). Lower boxes illustrate outcomes from agent POV. Upper circles and icons illustrate agent’s beliefs. Specifically, the lower box in (a) portrays the visual outcome of an empty horizon, and the auditory outcome of a soft sound. The upper portion of (a) provides some intuition as to the sort of alternative hypotheses that could be used to explain this sound. It could have been generated by a cat, a dog, or could be something else (e.g. an apple dropping from a tree). The last of these hypotheses corresponds to the belief that there is a small probability of hearing a sound, even in the absence of any creatures. The lower box in (b) shows the visual outcome of a dot on the horizon, in connection with the agent evaluating counter/actual beliefs (upper portion) that the visible dot could be caused by a distal cat or distal dog. (These scenarios recur in Figures 4 and 5.)

Scenario 2 (Figure 3b) also begins with Jerry roaming/scanning, and initially encountering a dot on the horizon. The dot can only be Tom or Spike, either approaching or retreating. Jerry will engage his defences if he infers Tom approaching, in order to keep Tom at a safe distance – as he strongly disprefers seeing Tom up close. But, he also disprefers being on the defence, which he infers via proprio- and baroreceptive cues. Alternatively, he could wait and see if Spike will appear upon closer examination, but this risks that it will turn out to be Tom after all. In other words, there is an irreducible ambiguity about whether the visible dot on the horizon is Tom or Spike, so Jeny must evaluate whether to await approach, orienting for more precise state estimation – i.e. exploration (epistemic foraging) – or to instead commit to defensive action, by affording greater precision to his prior preferences – i.e. exploitation (pragmatic foraging).

We now describe four variants for each of these scenarios. Jeny may believe that he is in a safe environment (’cat poor’) or an unsafe one (’cat rich’). This is reflected, respectively, by flat priors or by a strong prior that Tom is approaching. Another variation relates to the strength of the negative prior preference (i.e., aversion) for Tom. The hypothesis we test here is whether maiadaptive behaviour, such as that of animal and human PTSD, can be modelled by an exaggerated prior preference. To test this, we equipped Jeny with a negative prior preference for Tom that leads to emergent adaptive behaviour; we then strengthen this negative prior preference to simulate an ‘impaired’ Jeny, and demonstrate anticipated emergent maladaptive behaviour. Hence, the four variants for each of the two above scenarios are: (I) unimpaired / safe; (II) unimpaired / unsafe; (III) impaired / safe; and (IV) impaired / unsafe. A subset of the simulation results for two example trials for Scenario 1, Variant I are presented in Figure 4.

**Figure 4.**
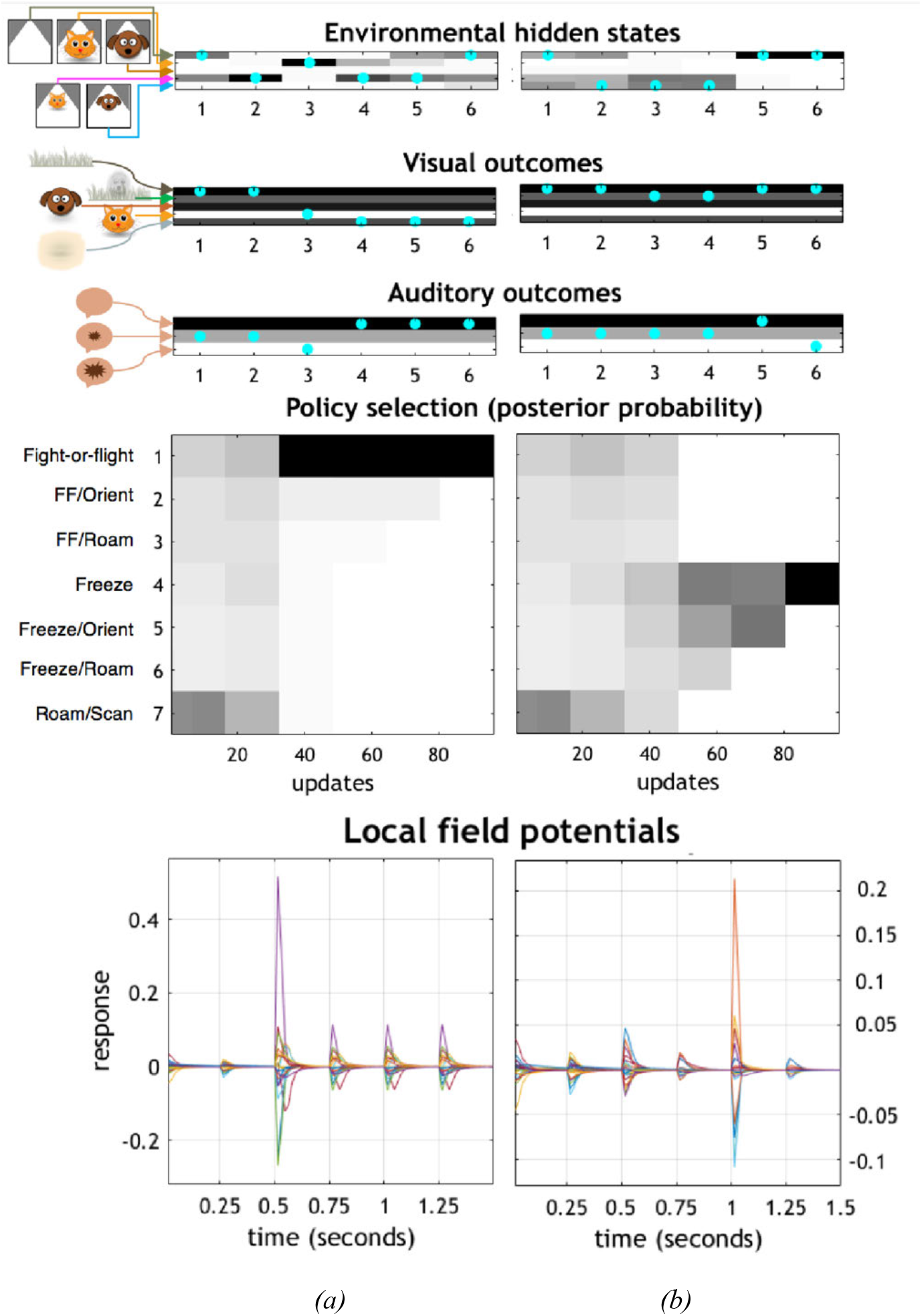
Simulated trials (columns a, b) for Scenario 1 (see Fig. 3a), Variant I (unimpaired/safe), illustrating the process of state estimation and policy selection. These plots show the trajectories of Jerry’s beliefs over the course of six time steps (x-axes) with five interleaved actions; where the action sequence comprises a policy. The grayscale spectrum in bars and upper box depict probabilities, from lightest (lowest probability) to darkest (highest probability). Cyan dots indicate the true state of the world (either states or outcomes. The shading in the outcome plots indicate prior probabilities (i.e. preferences), while those in the hidden state and policy plots indicate posterior beliefs. The graphics and text to the left of the plots show alternative states of the world and policies. Interpreting the posterior beliefs as neuronal firing rates, we can use their rates of change to generate synthetic local field potentials. Note that the increase in LFP amplitude coincides with the point at which Jerry becomes confident about the policy he is pursuing.

To connect this to the graphics in Figure 4, note that the first hidden state in Figure 4a (indicated by the cyan dot) shows no creatures in the vicinity. The outcomes generated by this state of affairs are an empty horizon and a soft sound (also indicated by cyan dots). At the second time-step, the cat is in the distance, and the same outcomes are generated. On Jerry’s next action – orienting toward the sound – he sees a cat shape and hears a loud sound, as Tom has by now entered a proximal range, which Jerry (correctly) infers. This inference induces a fight-or-flight policy selection, and rapid belief updating for policies as well as states averaged under policies. As can be seen in the LFP plot, this vigorous belief-update leads to a high amplitude LFP. The selected policy results in an increase in the distance between Tom and Jerry; hence the transition from a near cat to a distant cat from the third to fourth time-steps.

The sequence in Figure 4b shows that the dog enters distal range at the second time-step, causing a soft sound, but Jerry sees only an empty horizon. After Jerry orients toward the sound, he sees a distant dot on the horizon. However, at a distance, Jerry is unable to disambiguate between the hypotheses that the dot is caused by a cat or a dog. He has gone from wondering whether there is a cat, a dog, or nothing of interest (time-steps 1 and 2), to wondering whether there is a cat or dog (3 and 4). Only at the fifth time-step, when the dot recedes into the background, does Jerry attain precise beliefs about both the absence of a nearby creature, and about his ‘better safe than sorry’ freeze policy (as he has ruled out returning to a relaxed policy; see Figure 2). This causes the high amplitude LFP plotted above (Figure 4b). Please see the discussion for a more in-depth analysis of this scenario.

The distributions of policy selections for each variant over 32 trials per three conditions of Scenario 1 (4 × 32 × 3) and two conditions for Scenario 2 (4 × 32 × 2) are depicted in Figure 5. Note that either increasing aversion for the cat (impaired conditions), or specifying a prior belief that the cat is more probable (unsafe conditions), favour more frequent selection of more defensive policy choices, despite no change in the (hidden) states generating data. Having illustrated the characteristic behavioural responses that emerge under active inference, we now discuss how these behaviours are underwritten by emotional inference – and to what extent they can be considered a formal account of PTSD.

**Figure 5(a).**
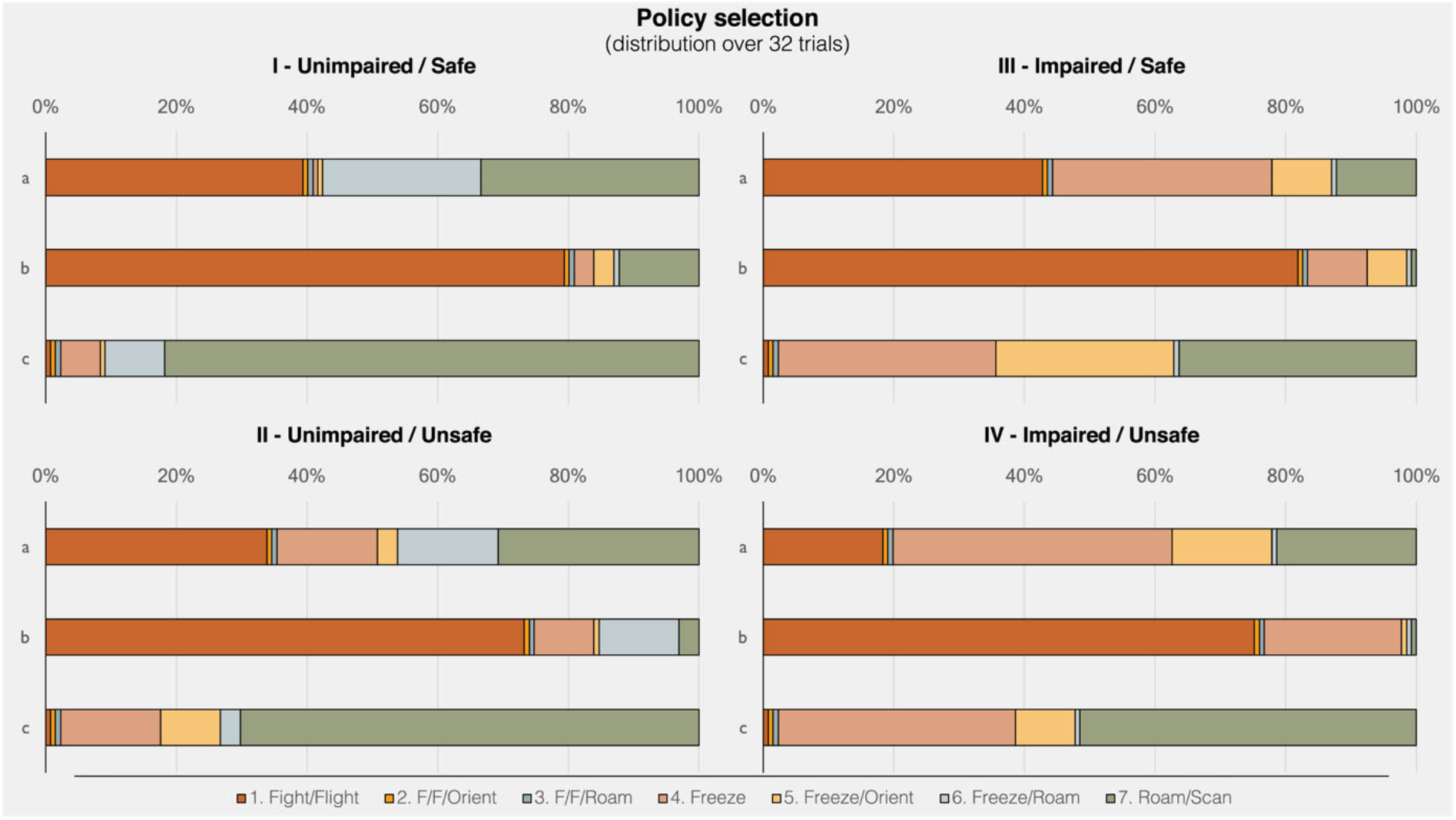
This figure reports the effect of perturbing prior beliefs on the sorts of actions selected. This is shown for a range of alternative initial hidden states. This action selection distribution is constructed from 32 trials of four simulated variants (I-IV) for Scenario 1 (Initial outcomes: empty horizon visible, soft sound audible – see Fig. 3a). Initial hidden state: (a) No creatures in the vicinity; (b) Distal cat; (c) Distal dog. Each bar depicts all policies in fixed ascending order (left to right), colour-coded (see key below bars), to aid comparison across bars of the relative increase or decrease in policy selection frequency. The variants serve to reveal the key influences on policy selection. I and II (unimpaired) are defined by a moderate prior preference; III and IV (impaired) are defined by an over-strong prior preference; I and III (safe) are defined by flat priors; and II and IV (unsafe) are defined by strong priors for a distal cat.

**Figure 5(b).**
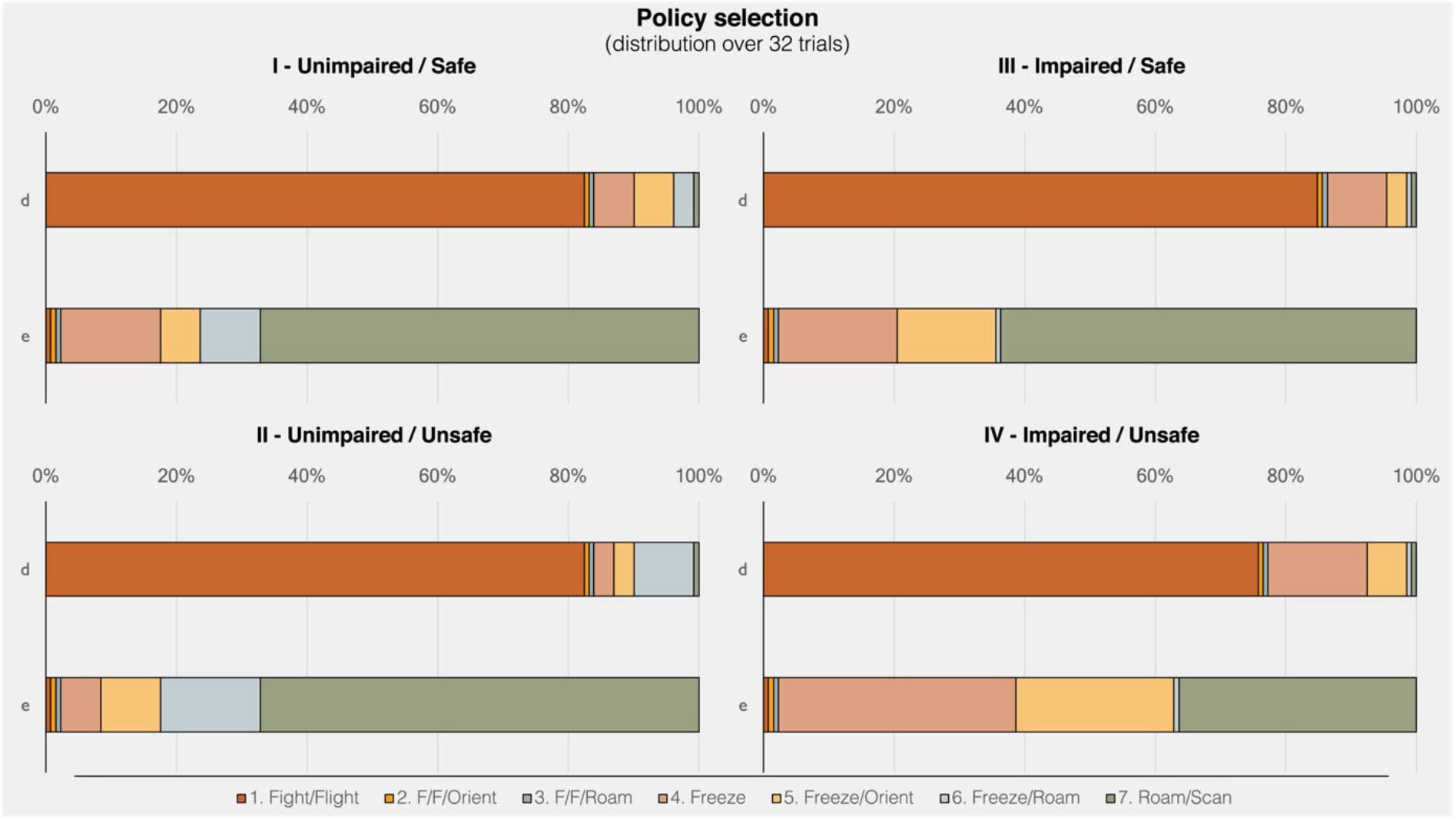
Policy selection distribution over 32 trials of four simulated variants (I-IV) for Scenario 2 (Initial outcome: dot on horizon visible – see Fig. 3b). Initial hidden state: (d) Distal cat; (e) Distal dog. (For further details, please see Fig. 5a caption and main text.)

## 4. Discussion

The normally intuitive relationship between ‘stress’ and ‘stressor’ – in the model on offer here – differs from conventional understanding. Under the generative model, causes are inferred from consequences, so an ordinarily un-stressful sensory outcome (e.g. a dot on the horizon or a soft sound), under some circumstances, can lead to the same inference as an overtly stressful cue (a close-up cat face); namely, clear evidence that a stressor is present. Given that actions affect hidden states, what is shown to be relevant above is that *stressor mitigation policies* may obtain even in the absence of stressful outcomes that lead to a stressor inference. If a stressor is a destructive force (state) with a high probability of challenging existential integrity (state transition to injury or death), for reasons of natural selection, there must be a precise prior aversion to its outcomes. Crucially, on this story, *any* outcome can induce a stressor mitigation policy selection, even in the absence of an actual stressor (i.e., false inference). From this perspective, the model offers a straightforward way to link a variety of animal and human models of PTSD, including predator, trauma, stress, and psychological and biological mechanism models (Matar et al., 2013).

### 4.1 Single trial phenomenology and neurobiology/electrophysiology

We can narrate the sequences depicted in Figure 4 in terms of Jerry’s phenomenology. Beginning with Figure 4a, by stipulation, the scenario begins while Jerry is roaming/scanning, and he is compelled to continue this first action until the second time-step. In the initial state, no other creatures are in the vicinity, such that Jerry sees only an empty horizon, but hears a soft sound. In the midst of roaming/scanning, he infers that the sound may be nothing interesting (e.g. a tree falling), or may be Tom approaching, so he seriously entertains continuing to roam/scan in a carefree manner. Meanwhile, his movement is in fact attracting Tom. By the second time-step, Jerry has become certain Tom is in the distance, even though he has not yet spotted his feline foe. This leads Jerry to a second action, orienting, which prepares a defensive response by raising his heart rate (not depicted) and also locates Tom in his visual field. By the third time-step, Jerry is both certain that Tom is too close, and certain that fight-or-flight is the most apt policy (actions 3-5).

In the lower portion of Figure 4a, we see a large LFP spike (after 500 ms.), with sufficient certainty in state estimation and defensive policy selection, and smaller spikes during subsequent state ambiguity. The relatively precise (posterior) beliefs about states and policies correspond to movement vigour, which would plausibly correlate with the selected defensive fight-or-flight response.

The sequence in Figure 4b has an identical set up, but reveals a different narrative and phenomenology. Under the same conditions, seeing an empty horizon and hearing a soft sound (with no creatures actually in the vicinity), Jerry considers the possibility that the sound may be nothing salient, or may be Tom *or* Spike distally approaching. Under the same constraints as above (roaming/scanning for action 1), at the second time-step, Jerry is still wondering whether the sound is nothing, Tom, or Spike. (In fact, Spike has wandered into the distal scene, out of view.) Jerry then orients toward the dot on the horizon (action 2), increasing his heart rate (not depicted), but this does not attract the approach of Spike, who has no interest in Jerry. In this case, Spike stays put, which keeps Tom away, but as Jerry cannot see who is in the distance, he remains concerned that it might be Tom. Thus, for action 3, Jerry holds still but has not yet fully committed to a freeze policy; he continues to entertain the possibility that it might be Spike in the distance after all, in which case he could resume carefree roaming/scanning. After holding still, whoever was in the distance (it happened to be Spike) has retreated rather than approached. Jerry is now sure no one is there, and he is correct. However, he decides roaming/scanning is not apt. He instead commits to a freeze policy (action 4-5), believing that this will ensure that Tom will not approach (better safe than sorry).

To track the above dynamics in the lower boxes of Figure 4b, we can observe a small LFP spike (after 500 ms.), with moderate certainty about a distal agent, and a large spike (after 1 second) at the elimination of remaining counterfactual policy selections – that would permit Jerry’s return to his preferred relaxed outcomes. This is consistent with studies of observed behaviour in non-human mammals, where a startle response proper – e.g., an abrupt, significant heart rate increase to a sudden, intense sound, with an accompanying behavioral shift to aroused and attentive, and reduced locomotor activity – is thought to be a precursor to a subsequent relaxation, fight-or-flight, or full-fledged parasympathetic freeze response (Lanier et al., 2000; Talling et al., 1996). Based on the present simulation, this division into sub-sequences could be regarded as a separation between an ‘a priori freeze’, continuous with orienting; during which further state estimation and/or policy evaluation is performed, followed by a defensive response proper, in this case, an ‘*a posteriori* freeze’. This analysis suggests an interesting consequence, namely, that a behaviourally continuous freeze state may be able to be empirically sub-divided into two phases with different underlying neurobiological and electrophysiological correlates.

### 4.2 Toward an alternative aetiology of psychological trauma: Reconceptualising fear and associative memory

With the present model, inferring a stress-related state, based on any outcome, entails inferring a policy that is expected to result in a transition to a stressor-free state. From first principles, we get three options that ‘fall out’ of the imperative to maintain my existential integrity by increasing the distance between me and the (inferred) danger: flight = get *farther* from danger; fight = get *further* from danger; freeze = get danger *farther* from me. Fear can therefore be regarded as a *post-hoc* construct for describing a phenomenon that emerges from those sensory outcomes which drive defensive action (in the service of self-maintenance), adaptively or maladaptively. This is apparent in the parallels between low-level, evolutionarily ancient neuronal survival architectures (e.g. in insects, for which a ‘fear emotion’ is less commonly referred to as such), and the more advanced higher-order human neuropsychology of trauma.

From this perspective, traumatic experience can be equated with an ecologically situated, embodied state that evinces high uncertainty in the sense of high complexity and low accuracy: many sensory outcomes are present and state estimation is confounded. During the trauma – on this story – a precise negative preference is set for an arbitrary outcome, whether or not it is in fact caused by the inflicting force; e.g. an intensely loud sound. This reframes ‘semantic’ associative memory as hypothetical inference; i.e. potentially false inference. For instance, instead of a loud sound being merely associated with an infliction of trauma, it becomes *evidence* for this, even when trauma will not in fact be inflicted. To revisit a common example, on the present account, it can be argued that it is not the car backfiring ‘cue’ that reminds the combat veteran of a gunshot by traumatic memory association – as it is characterised even in the otherwise comprehensive psychobiological literature (Pitman et al., 2012) – rather, the sound of the car backfiring is never inferred as being caused by a car in the first place. Instead, it is falsely inferred as caused by a gunshot, due to a learned hidden state-sensory outcome contingency. The learned relationship could have just as easily been formed with a quiet click rather than a loud bang, or any lifespan learning that assigns (potentially aberrant) high precision to prior preference.

On an evolutionarily timescale, a ‘learned’ inheritance can manifest as a conserved prior aversion; e.g. for intense sound (which could be ‘unlearned’ during the individual lifespan). In terms of natural selection, a ‘better safe than sorry’ adaptation increases survival odds, and is relevant not only to subsequent reproduction but also to niche construction (Constant et al., 2018), which would conserve the adaptation (e.g., learning to avoid untraversed paths by inferring they are hostile territory, without confirmation). To take a related example, an unexpected intense sound would normally be attributed with high uncertainty to its specific source. However, an adaptation that treats loud sounds as grounds for inferring (simply) a stressor; i.e. a biologically destructive force, renders a precise inference, precipitating a vigorous avoidance response, ranging from startle to flight. Notably, this can have a maladaptive downside as well, from natural population ecology up through human social psychopathology, especially in PTSD (Zanette and Clinchy, 2017).

This is evident in more basic neural architectures, such as flying insects, for which darkening fluctuations in the visual system (i.e. sensory outcomes) are sufficient to initiate escape behaviour (Holmqvist and Srinivasan, 1991). An even more closely related example occurs in crickets that respond to (predatory) bat ultrasound, for which a mesothoracic interneuron (homologous to the human scapula-trapezius innervation site) initiates an avoidance behaviour – a burst of motor activity – when a spike-rate threshold is met; this is normally caused by a sufficiently intense acoustic perturbation (Hoy et al., 1989). As the authors point out, it is thus one in the same if the neuron is defined as a ‘bat-detector’ or a ‘command neuron’. In our simulations, based on minimising expected free energy, both are recast as tightly coupled sensory estimation and motor policy selection to jointly secure existential continuity. Put simply, there are predicted future sensory outcomes that mandate defensive actions, whose consequences are the preferred future sensory outcomes. When uncertainty is sufficiently resolved for action, we see a (simulated) high LFP spike (Figure 4), offering a clear empirical prediction of measurable neurobiology. That is, a high intensity sensory outcome is cast as strong evidence that facilitates low uncertainty (precise) state estimation, and consequently precise policy selection.

It is not intrinsically important whether the stimulus intensity is high or low, but rather, what matters is that it correlates with a precise state estimation that precipitates avoidance. This puts a different spin on why olfaction is such an effective modality with respect to detecting predators. Their scent (outcome) is unambiguous, so by strongly dispreferring it, there will always be a relatively sure bet that the predator (state) can be avoided. That is, natural selection conserves the precise negative preference for particular olfactory outcomes, although other trade-offs come into play with purely chemosensory assessment (Kats and Dill, 1998). This can be contrasted with vision, which has greater ambiguity – you might mistakenly stay put and become lunch for a camouflaged or distant, fast-approaching predator – and auditory intensity, which has even greater ambiguity, given harmless loud events and dangerous quiet ones in the same frequency range.

In this light, from an information-theoretic perspective, in contrast to conventional animal communication and signalling theories, active inference allows for the emergence of a neuroethological and ecological story from first principles: it is a property of the generative model that multimodal integration lowers uncertainty or entropy. Empirical studies confirm that combining auditory and visual cues increases the robustness, precision, and discrimination of perceptual evidence relevant to decision-making (Best et al., 2007; Lewis and Noppeney, 2010; Noppeney et al., 2010). This is accounted for in the present simulations when combined auditory and visual cues resolve uncertainty about the presence of agents in the vicinity. Hence, natural selection favours multimodal integration, and conserves a ‘better safe than sorry’ fallback under greater uncertainty, such as when only audition or vision is available.

In short, depending on the neuronal architecture and character of the initial sensory evidence, agents bypass the question ‘what is it?’ and get straight to the point: ‘is there sufficient evidence that my integrity is in jeopardy?’. When the answer is ‘yes’, this implies evidence in favour of a pragmatic course of action – realising the preferred outcome by decreasing the sound intensity via fleeing – which will be a vigorous action in virtue of its relative certainty (Opris et al., 2011); i.e., a policy selection that minimises expected free energy by assigning high precision to prior preferences (Figure 4a). In this light, the adaptive/maladaptive and healthy/pathological distinctions relate simply to the sufficiency threshold of evidence that my integrity is in jeopardy. For some individuals, particular traumatic experiences would appear to semi-permanently lower this threshold such that it is easily met with imprecise evidence.

If the threshold is high enough, the intense auditory outcome might be initially met with a vigorous head turn, to perform quick epistemic foraging for uncertainty-resolving outcomes; for example, by foveating the source location (Heffner, 2004). The decision to select epistemic over pragmatic foraging – amounting to exploration outweighing exploitation – is based on an evaluation of counterfactual integrity-preserving actions. Namely, turning toward the source rather than fleeing immediately would conserve metabolic resources that would be wasted on an unnecessary diversion to skeletal muscles.^1^ Indeed, such unnecessary diversion takes place in maladaptive (unmerited) hyperarousal, discussed below.

### 4.3 Distributions of policy selections over 4 variants, 5 conditions, 32 trials per condition: Emergent hyperarousal and hypervigilance

Both the epistemic (orienting) and pragmatic (defensive) responses described above are consistent with the startle reflex, which can be contextualised within an ancient evolutionary sensorimotor architecture comprised of orienting and defensive vagal and/or sympathetic response mediation of the autonomic nervous system, especially the heart (Campbell et al., 1997). The exaggerated startle reflex and corresponding hyperarousal found in PTSD patients (Pole, 2007) is modelled here as unduly high precision on a prior preference that therefore more frequently evokes the pragmatic response. This emergent hyperarousal is reflected in Figure 5 as a decrease in light blue bands in all scenarios and conditions, and in Scenario 1 (Figure 5a) under presumed safe conditions (I and III) as a substantial decrease in green bands – and a corresponding substantial increase in light orange and light yellow bands. The impaired responses under presumed safety especially closely mimics PTSD-related hyperarousal, as the defensive policies obtain in situations for which the unimpaired agents frequently find no grounds for raising (or maintaining raised) defences.

A further nuance captured by this model relates to startled freezing behaviour. Consider that for an epistemically beneficial head turn (i.e., orienting), a possible consequence is the encounter of an aversive outcome and corresponding proximal threat inference, that in turn precipitates a pragmatic response (Friston et al., 2015a). In this case, vagal and/or parasympathetic lowering of the heart rate is likely invoked (‘freezing’). This action realises the pragmatic aim of maintaining a low probability of being detected by the threat (Roelofs, 2017), i.e. reducing the chance of a strongly dispreferred proximal threat outcome. However, it conflicts with the aim (preference) of avoiding vulnerable self-defensive state outcomes. When the latter aim wins out, heuristically, the stage is set for further epistemic foraging (Mirza et al., 2018).

The ability to transition out of a vulnerable self-defensive state therefore requires having low enough precision on the prior aversion, which can be interpreted as cognitive flexibility, associated with posttraumatic recovery (Yehuda and LeDoux, 2007; see also Sinapayen et al., 2017). An impaired agent – characterised here with excessively high precision on the prior aversion – may therefore become stuck in what closely mimics a hypervigilance behaviour (Figure 5b), a common PTSD symptom. Note that in presumed safety (I and III), and in the absence of actual danger (e), the contrast between the unimpaired (I) and impaired agent (III) is such that the former is able to resume roaming/scanning after a momentary freeze (light blue band) more frequently than the latter, who instead goes from freezing to aroused orienting (light yellow band). In presumed unsafe conditions (II and IV), in both actual danger (d) and its absence (e), the impaired agent (IV) also shows a substantial increase (approximately 4-fold) in unrelenting freezing (light orange band) compared to the unimpaired agent (II).

## 5. Summary and conclusions

We have presented a generative model of adaptive/healthy and maladaptive/pathological behaviour, grounded in an underlying evolutionary psychobiology. This computational approach offers a novel characterisation of PTSD based on first principles, rather than on conventional constructs such as ‘fear’ and ‘associative memory’, while remaining consistent with empirical findings linked to these constructs. The primary feature of the model is its use of overly precise prior preferences/aversions to model aberrant psychophysiological responses, where these precise priors are conserved by natural selection.

By treating the stressor (hidden state) inference as more fundamental than outcomes providing evidence for stress, we have proposed the following schema – that unifies standard active inference, neuroethological constructs, and a PTSD phenotype: namely, in specific contexts, exploration (epistemic foraging) is invoked for resolving uncertainty about the presence of a stressor. A high negative preference (set by evolution or learning) for a stressor-dependent outcome rapidly leads to exploitation (pragmatic foraging) before further state estimation. A ‘fearful’ or ‘aversive’ stimulus is thereby recast as an aversive outcome related to a stressor inference, with high precision assigned to the aversion.

Two common PTSD associated phenomena, the ‘generalisation of conditioned fear’ (a.k.a., an inability for disambiguation or discrimination of a fearful stimulus) and ‘fear extinction’ (a.k.a., safety learning) are also recast as two sides of the same coin – a dynamic balancing act between exploration and exploitation. Specifically, if excessively high precision is assigned to prior preference, this will limit the exploratory resolution of uncertainty regarding the inferred hidden state for a potentially stressor-dependent cue. Conversely, when this precision on preference is lowered sufficiently, the exploitative drive gives way to exploratory imperatives.

On this understanding, policies leading to a stressor-absent state are invoked for reducing uncertainty relative to precise prior preferences. These stressor-mitigation policies induce continuous arousal, and may invoke policies associated with movement vigour (e.g. fight-or-flight). Thus, startle is modelled as an aroused, vigorous response to a sensory outcome (e.g. sound), which may or may not transition into a defensive response, depending on a number of conditions. The conditions pertain to the fact that further exploration increases risk – that is, it raises the expected free energy of allowable policies because it lowers the probability of escaping the state that generates aversive outcomes. This points to a counterfactual policy evaluation delay, characterised here as an ‘a priori’ freeze, which differs from a defensive freeze, despite their behavioural identity. Moreover, this characterisation of startle permits us to model exaggerated startle, a common PTSD symptom, as arising from excessively precise prior preferences.

Finally, the model provides an explanation for the hyperarousal and hypervigilance found in PTSD patients. The former relates here to excessive (salience-related) motor policy readiness, due to such policies having the lowest expected free energy (lowest risk) for an active transition to a stressor-free state. This can emerge independently, or in conjunction with hypervigilance. The latter is framed as a sub-variety of excessive epistemic foraging, in which the predominant operative hypothesis (for which further evidence is sought) relates to the pragmatic avoidance of highly dispreferred outcomes.

In conclusion, active inference accounts for a wide range of evidence that, in some cases, has been disclosed within narrow disciplinary silos. The confinement of research to inherited constructs can at times suggest an impasse has been reached. However intuitive these predominant constructs may have become, rather than accept them *a priori*, we can reproduce construct-related phenomena as emergent from first principles. If, on this basis, synthetic PTSD can be shown to have adequate neuro-biobehavioural grounding – a project that will continue with future work – it should be possible to use this computational approach to build a bridge between basic research in psychobiology and translational therapeutic and clinical insights to improve health and wellbeing.

## Footnotes

1 Of course, if the embodied brain is there as a product of conserving metabolic reserves and the attending evolutionary imperatives, it need not know this; it must merely inherit precise (subpersonal) priors that amount to ‘I am not the sort of creature that moves unnecessarily’.

